# Assembly of gut-derived bacterial communities follows “early-bird” resource utilization dynamics

**DOI:** 10.1101/2023.01.13.523996

**Authors:** Andrés Aranda-Díaz, Lisa Willis, Taylor H. Nguyen, Po-Yi Ho, Jean Vila, Tani Thomsen, Taylor Chavez, Rose Yan, Feiqiao Brian Yu, Norma Neff, Alvaro Sanchez, Sylvie Estrela, Kerwyn Casey Huang

## Abstract

Diet can impact host health through changes to the gut microbiota, yet we lack mechanistic understanding linking nutrient availability and microbiota composition. Here, we use thousands of microbial communities cultured *in vitro* from human feces to uncover simple assembly rules and develop a predictive model of community composition upon addition of single nutrients from central carbon metabolism to a complex medium. Community membership was largely determined by the donor feces, whereas relative abundances were determined by the supplemental carbon source. The absolute abundance of most taxa was independent of the supplementing nutrient, due to the ability of fast-growing organisms to quickly exhaust their niche in the complex medium and then exploit and monopolize the supplemental carbon source. Relative abundances of dominant taxa could be predicted from the nutritional preferences and growth dynamics of species in isolation, and exceptions were consistent with strain-level variation in growth capabilities. Our study reveals that community assembly follows simple rules of nutrient utilization dynamics and provides a predictive framework for manipulating gut commensal communities through nutritional perturbations.

## Introduction

Microbial communities are critical components of practically every ecosystem on the planet (Berendsen et al., 2012; Falkowski et al., 2008; Human Microbiome Project, 2012), yet the nature of their assembly is largely mysterious. Key to their function is the establishment of a resilient community via interspecies and species-nutrient interactions that mediate community composition (Freilich et al., 2011; Harcombe, 2010; Momeni et al., 2013; Pacheco and Segre, 2019). The community within the mammalian gut constitutes one of the densest and most diverse bacterial ecosystems known (Backhed et al., 2005), and its composition is a strong signature of host health and disease. Gut commensals have diverse metabolic capabilities (Han et al., 2021) and diet is well known to alter gut microbiota composition (Backhed et al., 2004; David et al., 2014; Wu et al., 2011), suggesting that nutrients have a major impact on the metabolic functions of the gut microbiota. Yet, it remains unclear how nutrient availability and interspecies interactions, including nutrient competition (Goldford et al., 2018) and direct inhibitory interactions (e.g., via antibiotics, toxins (Stempler et al., 2017), and pH modification (Aranda-Diaz et al., 2020)) quantitatively contribute to the structure and function of gut microbial communities. Thus, the gut microbiota can provide a foundation for uncovering general rules of microbial community assembly and robustness, thereby revealing mechanisms for engineering communities with specific compositions and functions.

Phenomenological models focused on resource consumption (Ho et al., 2022b) have successfully predicted some aspects of microbiota variation, supporting a focus on metabolism for communities of gut commensals. Commensals exhibit a broad range of growth rates and levels of fastidiousness (Tramontano et al., 2018), which likely affects their ability to compete for nutrients, and differences in yield even among closely related strains (Tramontano *et al*., 2018) suggest distinct niches reflecting strain-specific differences in metabolic capabilities. Thus, many factors can potentially affect community assembly, and a major challenge is to identify growth variables that best predict composition and yield.

Mechanistic understanding of gut microbiota ecology and elucidation of the fundamental principles that underlie community composition and behavior have been hampered by low throughput and the fact that key parameters often cannot be precisely tuned in animal models (Fischbach, 2018). As a complementary approach, in recent years there has been increased interest in using *in vitro* culturing systems that afford better control of environmental parameters and increased throughput. Previous studies used *in vitro* co-culturing of collections of a small number of isolated species to probe interspecies interactions and metabolic functions (de Vos et al., 2017; Gutierrez and Garrido, 2019; Kehe et al., 2019; Venturelli et al., 2018), but these reconstituted communities have generally lacked the complexity of mammalian gut microbiotas. Another approach has focused on resuspension of stool in liquid media. These stool-derived *in vitro* communities (SICs) are highly stable and diverse, maintaining most major taxonomic families during passaging and after inoculation into ex-germ-free mice (Aranda-Diaz et al., 2022). Their composition varies across media (Javdan et al., 2020), and SICs have previously been used to study drug metabolism (Javdan *et al*., 2020; Zimmermann et al., 2019), as well as the response to antibiotics and resistance to pathogen colonization (Aranda-Diaz *et al*., 2022). Thus, SICs provide a useful model system in which to quantify the impact of nutrients on community assembly separate from complications of host metabolism, as well as whether assembly rules generalize across the broad range of compositions found in human hosts (Clemente et al., 2012).

Here, we show that SICs from human feces passaged in a complex medium supplemented with a single carbon source from central carbon metabolism exhibit inoculum-dependent membership (i.e., species presence/absence) but compositions (relative abundances) that depend on the supplemental carbon source. While the abundance of most families was approximately constant across environments, a few fast-growing taxa predictably increased in abundance with the supplemental carbon source. Focusing on a synthetic community of 15 strains isolated from a single parent community, we quantified the growth dynamics of these strains in isolation and in a reconstituted synthetic community and demonstrated that consumption of the supplemental carbon source largely follows depletion of individual niches in the complex medium. Iterative removal of the fastest-growing member typically promoted the second fastest-growing strain. Using a simple model that incorporates only isolate growth dynamics, we were able to predict the dominant members of a diverse range of communities. Moreover, the high degree of variability in family abundance across stool inocula in certain nutrient conditions was due to variation in the growth capabilities of the dominant strain in that family across donors. Taken together, this study suggests that community composition can be predicted based on species resource utilization dynamics across a broad range of environments.

## Results

### Stable SICs passaged from human stool inocula exhibit carbon source-dependent growth behaviors

To determine whether stool from different humans generate different *in vitro* communities, as we previously observed with humanized mice at different stages of antibiotic treatment (Aranda-Diaz *et al*., 2022), we passaged samples from three donors in BHI, one of the media that supported high diversity for humanized mouse SICs (Aranda-Diaz *et al*., 2022). Within <6 passages, SICs derived from human stool inocula converged to distinct, reproducible steady states with high diversity (~50 ASVs) that resembled the composition of the inocula (Fig. S1A,B). These data demonstrate that long-term batch culturing of human stool samples produces donor-specific, diverse, stable SICs, suggesting their potential for studying both general and specific ecological patterns across human microbiotas.

We sought to comprehensively characterize how nutrient availability affects human SICs via passaging on different carbon sources. We collected fecal samples from eight donors and inoculated them individually into 46 media (Fig. 1A,B) (Methods). Half of the growth conditions were minimal media composed of a base of M9 salts supplemented with one of 22 single carbon sources (sugars, alcohols, and short chain fatty acids). All carbon sources were provided at equimolar concentration to enable direct comparisons. The supplemental carbon sources spanned most of *Escherichia coli* central carbon metabolism, including the pentose phosphate pathway, glycolysis, and the tricarboxylic acid cycle (Fig. 1B). We also included M9 salts supplemented with no carbon source as a negative control for growth. The other 23 growth conditions were the same carbon sources along with a base composed of M9 salts and 10% BHI (henceforth referred to as BHI-based media), with the goal of increasing SIC diversity by providing a nutrient-rich base medium (10% BHI) that can support the growth of dozens of species (Fig. S1A) (Aranda-Diaz *et al*., 2022). We passaged these SICs in technical quadruplicate every 48 h for 7 passages to achieve stable compositions (Methods, Fig. S1A).

**Figure 1:**
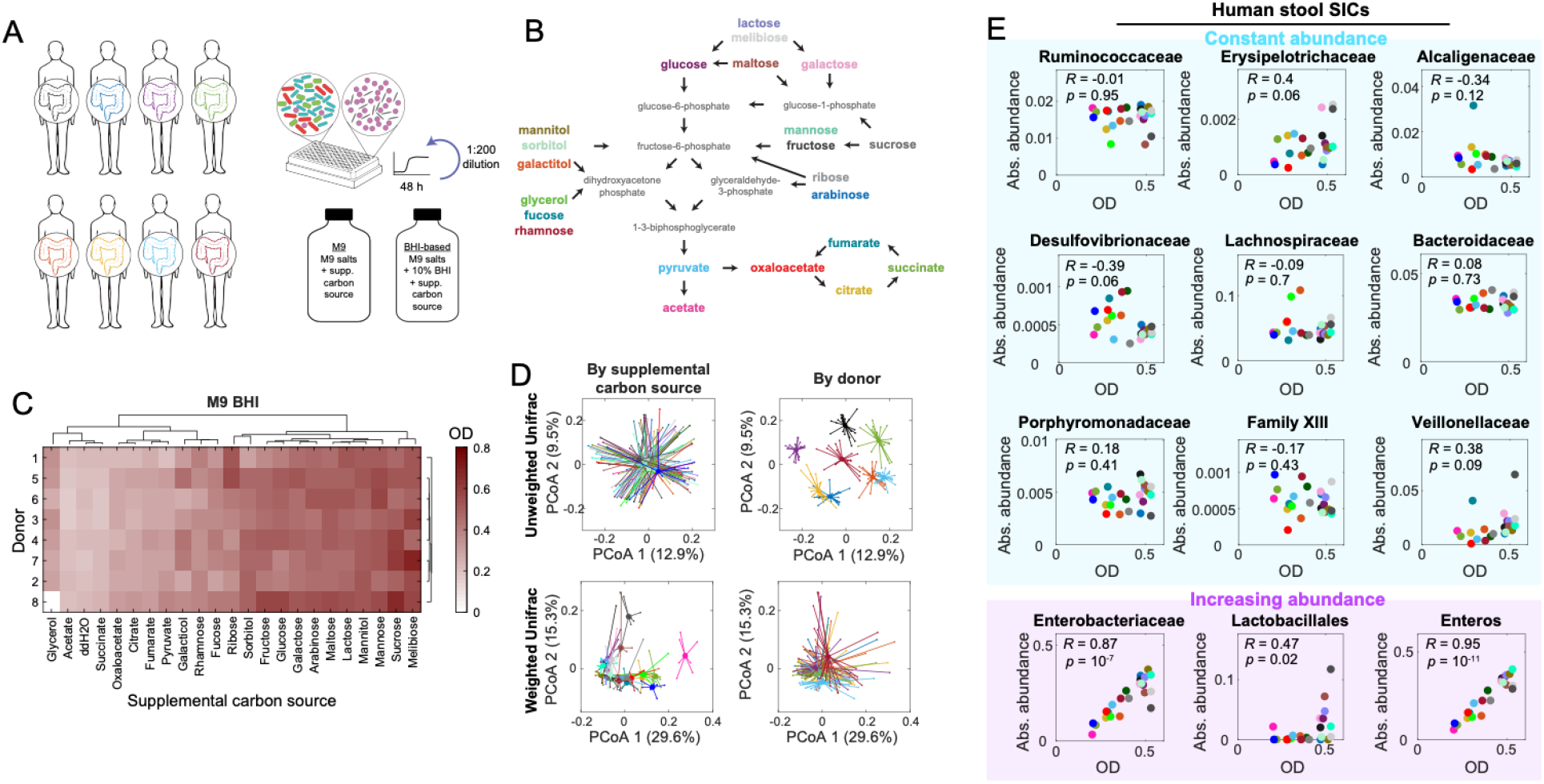
*In vitro* passaged communities separate by both carbon source and by donor. A) Stool from eight human donors was resuspended in 46 media (M9 salts without or with 10% BHI, supplemented with one of 22 carbon sources or ddH_2_O). The inocula were passaged in quadruplicate every 48 h for 14 days with 1:200 dilutions, resulting in 10,304 communities (8 donors × 46 media × 4 replicates × 7 passages) in total. B) The twenty-two selected carbon sources (bolded text) are metabolically diverse, including simple sugars, sugar alcohols, and short chain fatty acids. These carbon sources enter central carbon metabolism (CCM) at different points. C) The final optical density (OD) of the assembled communities (i.e., after the 7^th^ transfer) clustered approximately according to position in the CCM network. D) Top: using unweighted Unifrac (a metric that only considers taxa presence/absence, not abundance), community composition separated by donor and not by carbon source, indicating that the constituent ASVs were distinct across donors. Bottom: using weighted Unifrac, community composition separated by carbon source and not by donor, indicating that abundances are dictated by carbon source. Samples are colored by media (left) or donor (right). Circles represent the mean composition across replicates for each donor and carbon source, and are connected by lines to the centroid of all communities with the same carbon source (left) or donor (right). Here, the SICs from donor 1 were excluded from the weighted Unifrac analysis (Fig. S5C has PCoA plots with donor 1 included). E) Many families maintain a constant level across nutrient environments. Top (cyan): in M9+BHI-based human stool SICs, many families exhibited approximately constant absolute abundance (relative abundance multiplied by final OD) across BHI-based media supplemented with carbon sources despite wide variation in community yield. All *p*-values are >0.05. Bottom (magenta): the exceptions were the Enterobacteriaceae family and the Lactobacillales class (Streptococcaceae, Enterococcaceae, and Peptostreptococcaceae families), both of which increased in absolute abundance with increasing community yield. “Enteros” represents the sum of the Enterobacteriaceae and Lactobacillales absolute abundances, and was most highly correlated with community yield.

The SICs supported a broad range of yields with OD_final_~0-0.6, which were highly reproducible across replicates (Fig. S2A). To compare growth across SICs, we computed mean yield across the four replicates of each combination of donor and supplemental carbon source. We used hierarchical clustering of yield from the 7^th^ (final) passage separately for M9- and BHI-based conditions to determine if certain supplemental carbon sources consistently produced similar yields despite differences in inoculum composition. We identified approximately the same two clusters of supplemental carbon sources for M9 and BHI-based conditions, suggesting that the supplemental carbon source was a substantial determinant of community yield (Fig. 1C, S2B). In BHI-based SICs, sub-clusters were apparent: supplemental carbon sources involved in the tricarboxylic acid cycle clustered together, with acetate and succinate displaying the least growth. Another cluster of intermediate growth involved galactitol, fucose, rhamnose, and ribose, all of which are linked to pyruvate synthesis. The cluster with the highest yields involved the most upstream elements of central carbon metabolism, including pentose phosphate pathway compounds such as glucose, lactose, sucrose, mannitol, and sorbitol, in reasonable agreement with predictions from a flux-balance analysis (FBA) model (Methods; Fig. S3). These data suggest that metabolism is a major determinant of SIC assembly, motivating us to decipher the contributions of each SIC member.

### SIC membership is inoculum-dependent while relative abundances depend on supplemental carbon source

To determine whether differences in final yield were mirrored by compositional differences across supplemental carbon sources, we performed 16S sequencing of the SICs from their final passage to measure relative abundances. We then performed principal coordinate analyses (PCoA) of all combinations of 8 donors and 46 nutrient conditions. The M9 SICs clearly separated from the BHI-based SICs using weighted Unifrac (which incorporates taxon abundance as well as phylogeny) as a distance metric for the vector of relative abundances in each sample (Fig. S4A). Although some M9 SICs had >15 ASVs and up to 10 families present at >0.1% after 7 passages, their diversity was uniformly lower than all BHI-based SICs (Fig. S4B). These data show that SIC diversity is linked to medium complexity, as expected.

To interrogate the contributions of individual taxa to SIC yield in the context of communities, we focused on the more diverse BHI-based SICs. The two main modes through which we evaluated community diversity were membership (species presence/absence) and composition (relative abundance). To evaluate the former, in a PCoA using unweighted Unifrac (which incorporates phylogeny but not abundance) as the distance metric, SICs clearly separated by donor compared with supplemental carbon source (Fig. 1D), indicating that initial inoculum supercedes environmental conditions in determining SIC membership. Certain families drove the differences between donors (Fig. S5A,B). By contrast, in a PCoA using weighted Unifrac (which incorporates both phylogeny and relative abundance), the SICs derived from all donors except donor 1 were reasonably overlapping (Fig. S5C); donor 1 SICs had substantially lower levels of Enterobacteriaceae and substantially higher Veillonaceae and Enterococcaeae levels than any other donor (Fig. S5B) and thus were not included in subsequent PCoAs to enable cluster separation at higher resolution. Instead, the communities clustered by supplemental carbon source (Fig. 1D) and revealed an association between composition and yield (Fig. S6), suggesting that the relative abundance of SIC members is generally dictated by the supplemental carbon source, which affects community yield (Fig. 1C, S6). Taken together, the growth properties of these SICs demonstrate that presence or absence of SIC members and nutrient availability both drive SIC composition, and provide the potential to link nutrients, yield, and composition.

### The abundance of most low-abundance families is unaffected by supplemental carbon sources

To quantify the extent to which the supplemental carbon source dictates the abundance of each taxonomic group, we analyzed family-level relative abundances across all SICs after their final passage. Although SICs from the eight stool inocula exhibited a wide range of microbiota compositions, Enterobacteriaceae was the dominant family across all SICs (Fig. S5B), except for those from donor 1, which generally had very low levels of Enterobacteriaceae (Fig. S5B). The relative abundance of Enterobacteriaceae (the major family from the Proteobacteria phylum, which includes known generalists like the *Escherichia* genus that thrive on simple sugars) was similar across SICs from the other inocula on most supplemental carbon sources and was positively correlated with SIC yield across supplemental carbon sources (Fig. S7).

Other prevalent families exhibited relative abundances that were strongly negatively correlated with yield (Fig. S7). One possibility was that decreases in the relative abundance of certain families were a product of increases in the absolute abundance of other, high-abundance families like the Enterobacteriaceae. To address this question, we used the product of relative abundance with SIC yield (OD_final_) as a proxy for absolute abundance (Fig. 1E). With a few exceptions, Enterobacteriaceae absolute abundance increased linearly with SIC yield. Families in the order Lactobacillales (Streptococcaceae, Enterococcaceae, and Peptostreptococcaceae) also exhibited high abundance in some SICs with high yield (Fig. 1E). The summed abundance of Lactobacillales and Enterobacteriaceae (labelled as “Enteros”) accounted for almost all of the SIC yield boost from the supplemental carbon sources (Fig. 1E). By contrast, many other prevalent families, despite typically remaining at low abundance (<0.05), exhibited absolute abundances that were mostly independent of SIC yield across donors and were tightly distributed (Fig. 1E). Notably, noise in absolute abundance did not seem to be due to low relative abundances, since the Verrucomicrobiaceae family exhibited a tight distribution despite levels <10^-3^ (Fig. S7, S8). Moreover, the relatively constant abundance of many families across the 22 carbon sources suggests that metabolic networks and interspecies interactions are not strongly perturbed by the addition of simple carbon sources. Together, these data indicate that most of the supplemental carbon source is utilized by members of the Enterobacteriaceae or Lactobacillales and that the rest of the SIC members typically are unable to access the supplemental resources in the context of the community.

To evaluate whether direct passaging from feces into the various media was necessary for the supplemental carbon source to be monopolized by a few families without affecting the rest of the SIC, we studied the nutrient dependence of a stable SIC derived from a humanized mouse in 100% BHI (henceforth referred to as the “parent hmSIC”) (Aranda-Diaz *et al*., 2022). We regrew the stabilized parent hmSIC from a frozen stock in BHI and then passaged it twice in the 23 BHI-based media. The parent hmSIC rapidly transitioned to new compositions that were in quantitative agreement with SICs from human stool, with increased absolute abundances of the Enterobacteriaceae and the Lactobacillales that were correlated with yield and almost no changes in the other families (Fig. S9). These data show that SICs rapidly transition to a steady state in which a few families monopolize resources, and that the taxonomy of the strains that monopolize those resources is conserved across communities.

### Supplemental nutrients affect isolate growth dynamics only after consumption of preferred resources in BHI

One possible explanation for the increased abundance of the Enterobacteriaceae and Lactobacillales while the other families maintained constant abundance across many supplemental carbon sources is that the Enterobacteriaceae and Lactobacillales are the only species able to consume the supplemental carbon source. To test this hypothesis, we used 15 strains isolated from the parent hmSIC (Methods), representing 6 families and comprising 69% of the total abundance of the parent hmSIC (Fig. 2A). We screened each of the 15 isolates and laboratory *E*. *coli* strain MG1655 (an Enterobacteriaceae) in the 23 BHI-based media. Contrary to our hypothesis, we found that many strains are generalists and gain a growth advantage from the additional carbon sources. Clustering based on isolate yield across media was largely consistent with phylogeny. The two Enterobacteriaceae species, *E. coli* and *Escherichia fergusonii*, exhibited some of the highest yields on most supplemental carbon sources (Fig. 2B), except on acetate, citrate, succinate, and sucrose, consistent with the observation that the latter set were supplemental carbon sources on which SICs exhibited relatively lower abundances of Enterobacteriacae (Fig. 1E, S9). Nonetheless, there were many strains that could utilize most carbon sources, often with high yields. *Enterococcus* species had high yields in sucrose (Fig. 2B), a carbon source in which Lactobacillales dominated SICs (Fig. 1E, S9), but *B. fragilis* and *C. clostridioforme* also exhibited high isolate yields on a subset of carbon sources (Fig. 2B), despite being unable to compete for these nutrients in a community context. Thus, the ability to utilize a carbon source in isolation did not explain the general dominance of Enterobacteriaceae in SICs.

**Figure 2:**
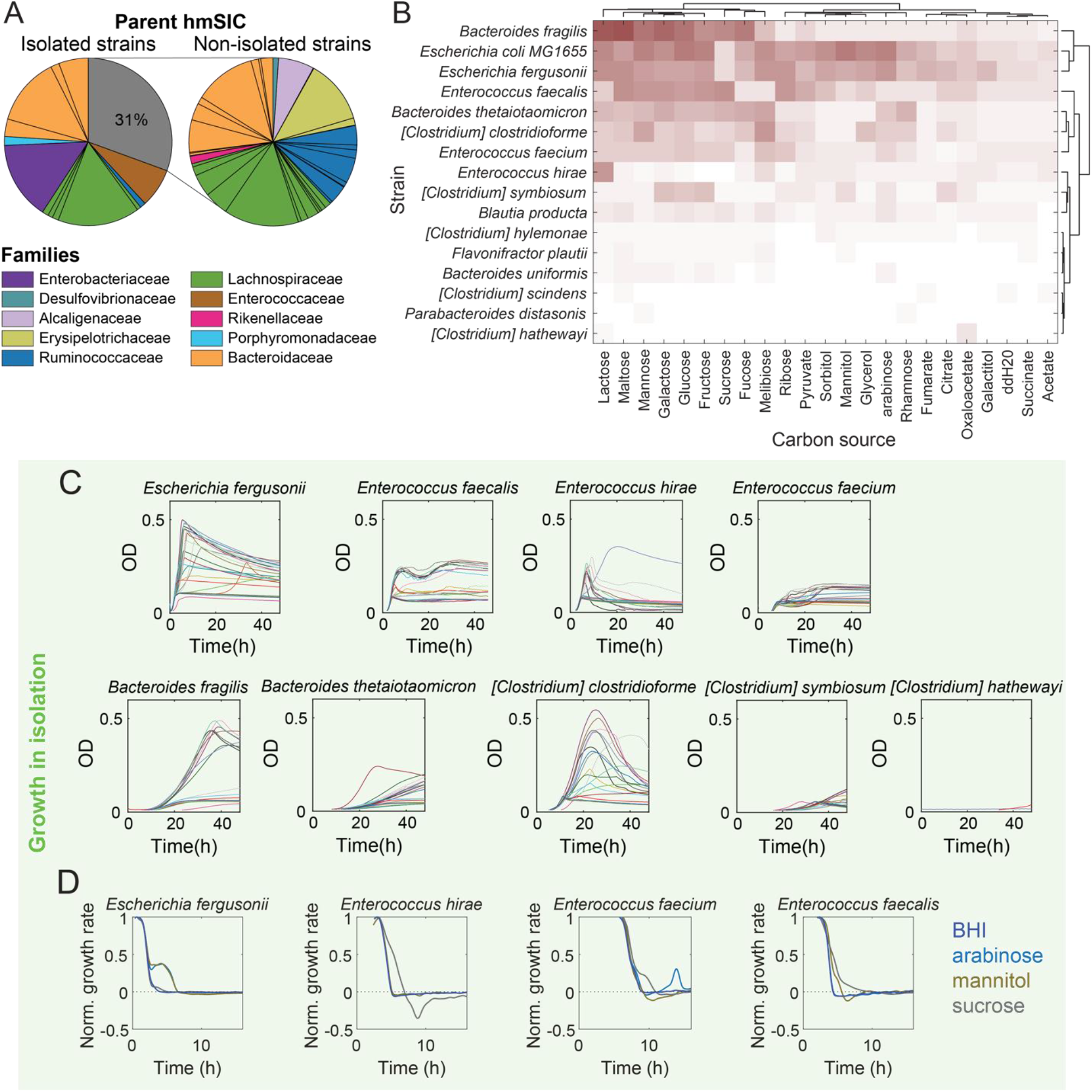
Individual species capitalize on nutrients from BHI before transitioning to the supplemental carbon source. A) Fifteen strains isolated from the hmSIC accounted for most of the families and 69% of the abundance. B) Yield for 15 isolates and *Escherichia coli* MG1655 in 10% BHI plus 22 carbon sources or ddH_2_O clustered approximately based on phylogeny. Many isolates were able to grow well in many carbon sources. Carbon sources clustered similarly as in the SICs derived from human donors (Fig. 1C). C) Growth curves of isolates were quantitatively similar across 10% BHI supplemented with one of 22 carbon sources or ddH_2_O for the initial period of growth, suggesting that the species first consumed nutrients in BHI and then switched to the supplemental carbon source. D) Growth rate normalized to its peak value was initially similar across carbon sources, after which each species transitioned to consumption of the carbon source with distinct dynamics. Arabinose, mannitol, and sucrose are highlighted as representative examples.

Interestingly, however, the supplemental carbon source tended to only affect isolate growth dynamics in the later stages of the growth passage. Indeed, for almost all strains and supplemental carbon source pairwise combinations, the initial phase of growth was virtually identical to that in base medium, up to the time when OD plateaued in base medium (Fig. 2C). Afterward, OD continued to increase for many strains when a supplemental carbon source was present (Fig. 2C-D, S10). This observation suggests that species first exhaust their niche on BHI and then switch to consume the additional carbon source, and more generally that the sequence of resource utilization, rather than simply the ability to consume resources may play an important role in shaping community composition.

### SIC members exhibit a wide range of growth dynamics in the base medium

The dominance of Enterobacteriaceae, some of which can be fast-growing (e.g., *Escherichia coli* (Atolia et al., 2020)), across many supplemental carbon sources suggested that the structure of a community might be dependent on the growth dynamics of each member. The maximum growth rate of the parent hmSIC in 10% BHI without any supplemental carbon sources (base medium) was attained in the first few hours and was equivalent to a doubling time of ~35 min (Fig. 3A, B), consistent with the hypothesis that a fast grower dominated early growth. Consistent with this idea, the presence or absence of a fast-growing species from the Enterobacteriaceae or Lactobacillales was predictive of maximum growth rate across a collection of SICs from humanized mice (Fig. S11).

**Figure 3:**
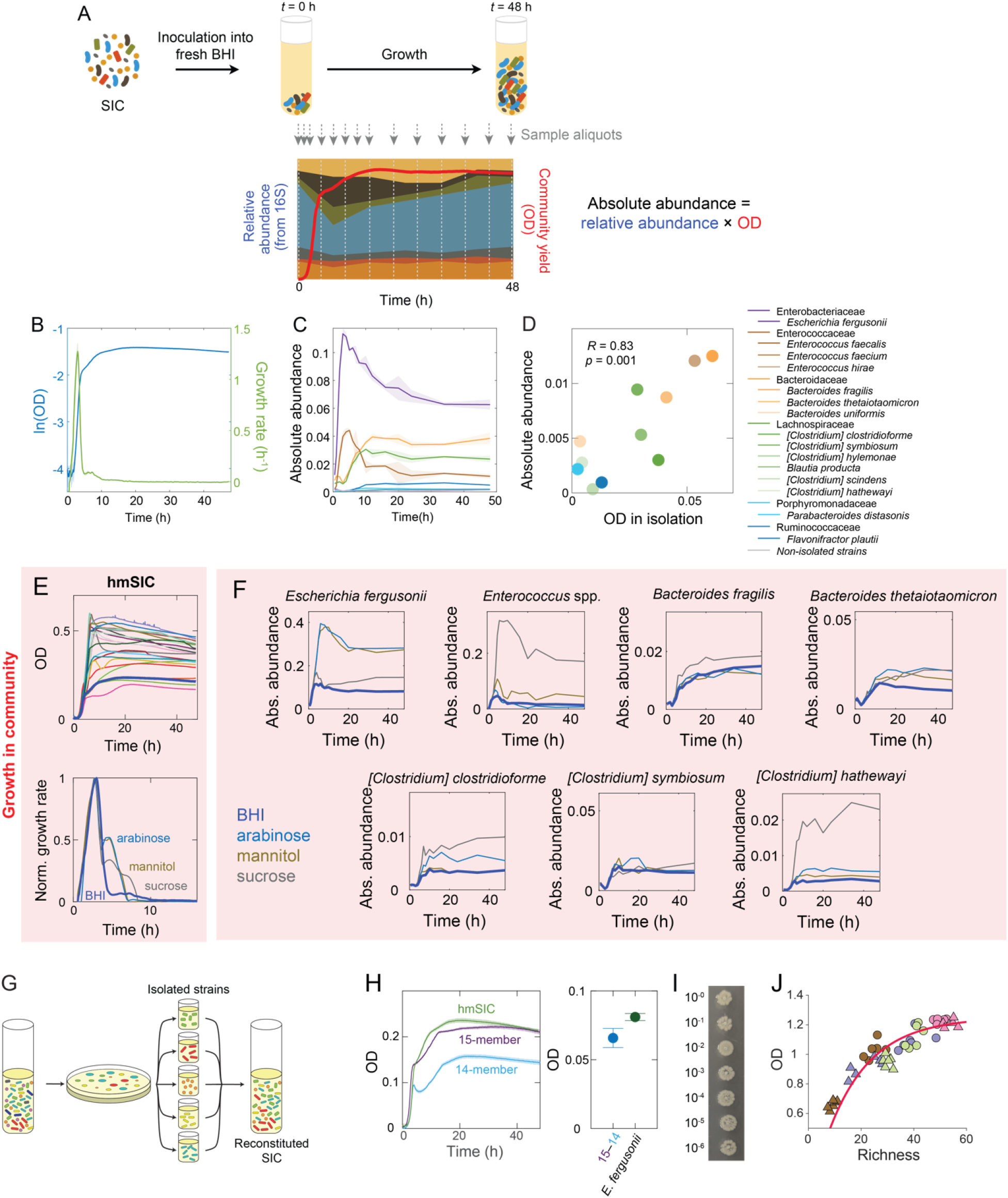
Community members display a wide range of growth behaviors in isolation that are correlated with abundances in a community context. A) The abundance dynamics of each species in a community were tracked by sampling an aliquot at various time points during a single passage (48 h of incubation) in BHI. 16S rRNA gene sequencing of the aliquots was used to assess the relative abundance of each species in the community. Community yield (OD) was measured concurrently. The absolute abundance of species *i* at each time *t* was calculated by multiplying its relative abundance by OD. B) Growth rate (green) of the hmSIC during growth in BHI as computed from OD (blue) peaked after a few hours and saturated after ~10 h. C) The absolute abundance dynamics of each family within the hmSIC was computed as shown in (A). Enterobacteriaceae and Enterococcaceae levels increased more rapidly than all other families. D) For the 15 isolates, final OD when grown in isolation was correlated with absolute abundance in the hmSIC. E) Top: growth curves of the hmSIC across BHI-based media. Bottom: similar to isolates (Fig. 2D), normalized growth rate was initially similar across carbon sources before a transition to carbon source consumption. F) Absolute abundance dynamics of individual species within the hmSIC estimated from sampling as in (A) supports the hypothesis that each species first consumes nutrients within its niche in BHI. *E. fergusonii* and *Enterococcus* species then transitioned to consuming the supplemental carbon source, while most other species displayed little growth on the supplemental carbon source. Bacteroidaceae and Lachnospiraceae species grew more quickly in a community than in isolation (Fig. 2C), suggesting cross-feeding from Enterobacteriaceae and/or Enterococcaceae members. G) Versions of the 15-member SIC were reconstituted with strains isolated from the hmSIC (“reconstituted SIC”). H) Left: the 15-member reconstituted SIC reached a similar final OD to the hmSIC when grown in 10% BHI, whereas the 14-member SIC lacking *E. fergusonii* had significantly lower yield. Right: the difference in final yield between the 15- and 14-member SICs was similar to the yield of *E. fergusonii* when grown in isolation in 10% BHI. I) In a 15-member community reconstituted with equal OD inocula of every species except *E. fergusonii*, the abundance of *E. fergusonii* after 1 passage in 100% BHI was approximately independent of its initial density. Dilution levels shown are relative to a 1:200 dilution of a stationary-phase *E. fergusonii* culture. J) Across SICs derived from humanized mice at various time points before and after treatment with ciprofloxacin, richness (number of ASVs) was highly correlated with final OD across communities. A consumer-resource model successfully predicted this relationship (pink), with a sparsity factor (probability that an ASV cannot utilize a given resource) of *S*=0.94.

To test whether the dynamics of species in a community were reflective of their growth in isolation and further dissect the dynamics of each member within the parent hmSIC, we passaged the parent hmSIC in base medium and sampled an aliquot for 16S rRNA gene sequencing every 30 min or 1 h for the first 8 h and every 2-16 h thereafter (Fig. 3A, Methods). We also monitored OD to estimate absolute abundances from 16S relative abundances throughout the growth curve. The Enterobacteriaceae and Enterococcaceae saturated their growth in BHI at least 5 h before any other families (Fig. 3C), as expected since most of their members are fast growers. The Lachnospiraceae and Bacteroidaceae showed significant growth in the interval between ~3-10 h. Individual members within those families showed a variety of growth dynamics, with certain species like *Clostridium clostridioforme* and *Bacteroides fragilis* growing to high levels within 12 h (Fig. S12). These data indicate that the abundance of some members will saturate in base medium long before others.

While the parent hmSIC had a higher yield than any of the isolates, isolates typically had lower estimated absolute abundances in the SIC compared to their yield when grown alone (Fig. 3D), suggesting that they experience competition for resources in BHI. Nonetheless, absolute abundance in the parent hmSIC was correlated with isolate yield (*R*=0.83, *p*<0.001, Fig. 3D) but not with maximum growth rate (Fig. S13A) or lag time (Fig. S13B). The observation that even isolates with very low maximum growth rate (e.g., *Clostridium hylemonae*) can achieve abundances within a community commensurate with their yield in isolation suggests that, for many strains, a subset of the nutrients that they consume are distinct from the other strains. Furthermore, for some members such as the Bacteroidaceae, growth was faster in the parent hmSIC (Fig. 3F) than in isolation (Fig. 2C), suggesting that metabolic cross-feeding enhances growth in the parent hmSIC and obscures potential correlations between relative abundance and maximum growth rate/lag time in isolation. Thus, parent hmSIC assembly in BHI is a combination of growth partitioned into specialized niches, resource competition, and cross-feeding.

### A second phase of growth underlies the ability to access supplemental carbon sources

Similar to the isolates, the parent hmSIC also showed a second, slower phase of growth that led to an increase in yield relative to BHI in most supplemental carbon sources (Fig. 3E). To examine how community dynamics were related to the absolute abundances of each constituent strain, we passaged the parent hmSIC in all 23 BHI-based media and sampled an aliquot for 16S sequencing every 30 min or 1 h for the first 8 h and every 2-16 h thereafter (Fig. 3A). Regardless of the supplemental carbon source, the initial dynamics of each species in the first few hours were essentially the same (Fig. 3F). *E. fergusonii* and *Enterococcus* species expanded with dynamics similar to their growth in isolation (Fig. 2C, 3F), while *Bacteroides* and *Clostridium* species increased on a shorter time scale than in isolation due to a combination of shorter lag and higher growth rate (Fig. 2C, 3F), again suggesting fitness benefits from Enterobacteriaceae and/or Enterococcaceae growth. At or shortly after the time that the Enterobacteriaceae and Enterococcaceae growth would have saturated in base medium, they increased in absolute abundance in the hmSIC in a second wave of growth on the supplemental carbon source (Fig. 3F), consistent with their growth in isolation (Fig. 2C, 2D). These data suggest that the growth capabilities of fast-growing strains in isolation and their competition for additional resources in SICs drive growth dynamics and SIC composition when a complex base medium is supplemented with a simple carbon source.

### The abundance of a fast-growing member is independent of its initial density

When grown as a community, if fast-growing strains are able to use their preferred resources as well as the preferred resources of other strains faster than slow-growing strains, one would expect fast-growing strains to increase in relative abundance over the course of repeated passaging in fresh medium. However, the observation that all strains, including *E. fergusonii*, exhibit constant absolute abundance within SICs over multiple passages suggests that BHI is stratified into well-defined nutrient utilization profiles such that each strain mainly consumes certain resources either uniquely or faster than all other strains.

To test this hypothesis, we sought to complement our top-down approach using SICs with bottom-up reconstitution using the 15 isolates (Fig. 3G). A co-culture of these 15 strains reproduced most of the growth properties and composition of the parent hmSIC in 100% BHI (Fig. S14). In base medium, the difference in final yield between the 15-member community and a 14-member community lacking *E*. *fergusonii* was quantitatively similar to the final yield of *E. fergusonii* in isolation (Fig. 3H), suggesting that a majority of *E. fergusonii’*s niche is distinct from that of the other strains.

Additionally, we reconstituted the 15-member community with an equal OD mixture of all strains with the exception of a dilution of up to 10^-6^ for *E. fergusonii*. A 10^-6^ dilution reduced the *E. fergusonii* inoculation size to <100 cells, and required ~20 additional generations of its growth to reach the same absolute abundance. This additional growth required ~8-10 h at *E. fergusonii’*s maximum growth rate, which would delay its saturation behind that of several other strains (Fig. 3C). Nonetheless, the abundance of *E. fergusonii* after a single passage was not substantially affected by dilution (Fig. 3I), indicating that the delay in growth due to severe dilution did not allow any other strains to take over a large fraction of *E. fergusonii’*s preferred nutrients.

### A consumer-resource model with sparse resource utilization predicts the scaling of community yield with diversity

To examine the significance of this nutrient partitioning, we took advantage of a set of SICs from our previous study that were derived from mice inoculated with the same human fecal sample (and hence involved the same pool of strains) and treated with antibiotics for different amounts of time before sampling feces to generate SICs (Aranda-Diaz *et al*., 2022). Across SICs passaged in BHI, yield varied over a wide range and was highly correlated with SIC richness (*R*=0.92, *p*=5×10^-20^; Fig. 3J). Thus, for a fixed nutrient supply, SIC yield is greater when more taxa are present, suggesting that the addition of more members leads to a larger number of consumed metabolites (although more taxa may also lead to more constructed niches and opportunities for cross-feeding (Estrela et al., 2022a)).

To determine whether such a relationship between richness and yield is expected to be a general property of SICs, we investigated a consumer-resource model with differently sized subsets of strains sampled from a fixed resource consumption matrix (Methods). Within this model, the relationship between richness and yield was qualitatively in agreement with our experimental data (Fig. 3J). The quantitative nature of this relationship was dependent on the probability that a strain does not consume a nutrient (*S*, the sparsity of the resource-consumption matrix). We fitted our experimental data at the ASV level and obtained a predicted sparsity of *S*~0.94 (Fig. 3J), meaning that an ASV on average is capable of consuming 6% of the available metabolites. Intriguingly, this estimate is in close agreement with metabolomics-based measurements of the metabolites consumed by the 15 species used in this study (Ho et al., 2022a). Thus, the application of metabolic modeling to a community exposed to a broad range of perturbations provides quantitative insight into the competition landscape.

### A simple model predicts community assembly based on isolate growth

Given that the dominant species in SICs were generalists and fast growers, we sought to understand the determinants of this oligodominance. To further explore the identity of the dominant strain (or ASV in the case of the parent hmSIC) and the extent to which they dominated supplemental carbon sources, we calculated each strain’s share of the excess yield in the community:

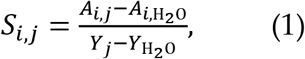

where *S_i,j_* is the fractional share of strain *i* in carbon source *j*, A_i,j_ and A_i,H_2_O_ are the absolute abundance of strain *i* in supplemental carbon source *j* or H_2_O (in addition to BHI), respectively, and *Y_j_* and Y_H_2_O_ are the yield of the community in supplemental carbon source *j* and H_2_O, respectively. Note that the sum of *A_i,j_* over all strains *i* in carbon source *j* is *Y_j_* based on our definition of absolute abundance. This metric showed that *E*. *fergusonii* accessed the majority fraction of the excess yield in many supplemental carbon sources, but also shared a fraction of the yield with *Enterococcus* species in several cases (Fig. S15). To test whether we could reconstruct the carbon source-dependent growth behaviors of the human donor-based SICs (Fig. 1) or the parent hmSIC (Fig. 3) in a defined (reconstituted) community, we grew a mixture of the 15 isolates discussed above for 1 passage in base medium (BHI only) to allow it to equilibrate, and then grew the resulting 15-member community in all 23 BHI-based nutrient conditions for 2 passages. The yield in most carbon sources was very similar to that of the parent hmSIC (Fig. S16A), suggesting that the 15-member community is able to make use of the available carbon to similar extent as full SICs despite missing 30% of the abundance (Fig. 2A), presumably because the most abundant families all have isolate representatives. The Enterobacteriaceae and Enterococcaceae dominated in most carbon sources (Fig. 4Aii) with similar patterns of coexistence as in the hmSIC (Fig. 4Ai) and most families maintained constant absolute abundance across carbon sources as in the hmSIC and human-derived SICs (Fig. 1E, S9). In general, carbon sources that were dominated by either Enterobacteriaceae or Enterococcaceae showed large differences in the ability and timing of utilization of those carbon sources (sucrose and arabinose) by these dominant strains, in isolation (Fig. 2B,C) and in SICs (Fig. 3F). The Enterobacteriaceae and Enterococcaceae were equally competitive in carbon sources for which their dynamics of carbon source utilization in isolation were similar (e.g., maltose; Fig. 2C). Other species such as *C. clostridioforme* and *B. fragilis* were capable of achieving high yields on those carbon sources in isolation but exhibited growth at later times than the Enterobacteriaceae and Enterococcaceae (Fig. 2B-2C), explaining their inability to dominate in communities. These observations led us to hypothesize that both the ability to utilize a supplemental carbon source and the timing at which utilization initiated after exhaustion of its BHI niche determine the extent to which a strain dominates.

**Figure 4:**
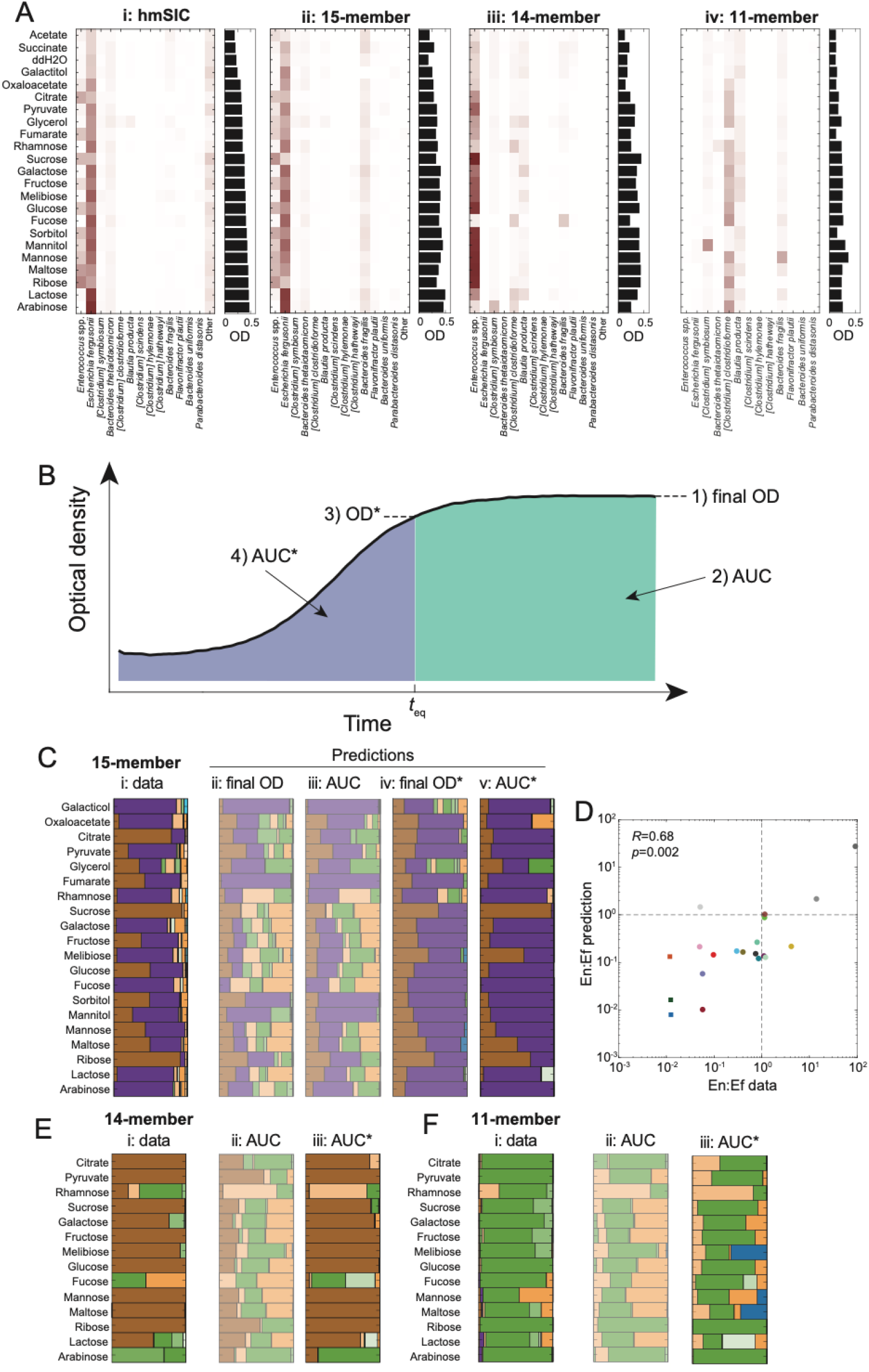
Isolate growth dynamics quantitatively predict capacity to access yield from supplemental carbon sources. A) The relative abundances of *E. fergusonii* and/or *Enterococcus* species mimicked that of the entire community (bars on the right) for (i) the hmSIC, (ii) the 15-member community, (iii) the 14-member community, and (iv) the 11-member community, with similar patterns in (i) and (ii). B) To quantitatively predict each strain’s share of the excess yield in the community beyond growth in the BHI-base medium alone, we used data from isolate growth curves and evaluated Eq. 1 using different growth parameters: 1) the final OD or 2) the area under the curve (AUC; blue + green regions) of the entire isolate growth curves; 3) the final OD (OD*) or 4) the AUC (AUC*; blue region) of the curves up to the time *t*_eq_ at which the sum of the OD of the isolates equaled the yield of the *N*-member reconstituted community (where *N*=15, 14, or 11). C) (i) The excess yield computed from Eq. 1 for the 15-member community exhibited similar patterns to the hmSIC (Fig. S16). Predictions of excess yield based on the final OD of isolate growth curves (ii) or area under the curve (iii) failed to predict excess yield. Final OD (OD*) at the time at which the summed excess yield of the 15 isolates equaled that of the community (*t*_eq_) predicted relatively equal dominance by *E. fergusonii* and *Enterococcus* species (iv), while the AUC at *t*_eq_ (AUC*) accurately predicted most compositions (v). Shown are all carbon sources for which the AUC* prediction can be robustly determined. D) Predictions of the ratio of *Enterococcus* spp. to *E. fergusonii* abundances (En:Ef) based on AUC* were highly correlated with experimental data. E,F) Predictions of composition based on AUC* (iii) more successfully recapitulated experimental data (i) than predictions based on the AUC of the entire isolate growth curves (ii), for both the 14-member community (E) and the 11-member community lacking *E. fergusonii* and all three *Enterococcus* spp. (F). Shown are all carbon sources for which the AUC* prediction can be robustly determined.

To test this hypothesis quantitatively, we evaluated several strategies to predict each strain’s share of excess yield using our data from growth curves in isolation (Fig. 4B). First, we subtracted the growth curve in base medium alone from growth in base medium with each supplemental carbon source and used these differences to predict the composition of excess yield (Fig. 4Ci). As expected, final OD (Fig. 4Cii) or the area under the curve (Fig. 4Ciii) of the difference of the two growth curves did not recapitulate experimental measurements of excess yield share; for instance, the share of late growers such as *B*. *fragilis* and *C*. *clostridioforme* was overestimated.

To account for the timing of carbon source utilization, we used data from isolate growth curves up to the time *t*_eq_ at which the sum of the OD of the isolates equaled the yield of the 15-member community in that carbon source (Fig. 4B), which we estimated to represent the time of resource exhaustion in the community. The OD of the isolates at that time point (“final OD*”) led to a reasonable prediction of the composition of the excess yield, although the abundances of *Enterococcus* strains were much more uniform than in experiments (Fig. 4Civ). The area under the curve up to the time *t*_eq_ (“AUC*”), which is influenced by both utilization ability and timing and hence links naturally to depletion of all nutrients utilized by the community, successfully predicted most of the compositions of excess growth across carbon sources (Fig. 4Cv), even the quantitative ratio of *Enterococcus* strains to *E. fergusonii* abundance (Fig. 4D).

### Elimination of the fastest grower primarily promotes another rapidly growing strain

We hypothesized that the accumulated growth of isolates up to the point when resources were utilized in the community would generally be a good predictor of community assembly with the supplemental carbon source. To test this hypothesis under conditions where the organism consuming the supplemental carbon would necessarily change, we grew a 14-member community lacking *E. fergusonii* and an 11-member community lacking *E. fergusonii* and all *Enterococcus* strains in most of the 23 BHI-based nutrient conditions. Yields of the 14-member community were similar to, albeit slightly lower than, that of the hmSIC (Fig. 4B, S16B), as expected given the absence of certain strains (Fig. 2A). Similar to the parent hmSIC and 15-member community, calculation of the expected share of excess growth in the 14-member community based on isolate growth correctly predicted the ability of the Enterococcaceae to expand in most carbon sources (Fig. 4C, 4Ei) when based on AUC up to *t*_eq_ (Fig. 4Eiii) but not when using the entire growth curve (Fig. 4Eii). Isolate growth predicted that Lachnospiraceae and Bacteroidaceae such as *C. clostridioforme* and *B. fragilis* would access a larger share in some carbon sources (rhamnose, fucose, lactose, and arabinose). These predictions were largely born out in 14-member communities, although the relative proportions of Lachnospiraceae and Bacteroidaceae were sometimes predicted poorly. Predictions of the composition of excess growth in an 11-member community lacking the Enterobacteriaceae and all Enterococcaceae largely matched experimental data (Fig. 4F), particularly the dominance of *C. clostridioforme*. In sum, isolate growth up to the point at which the supplemental carbon source is predicted to be fully depleted can predict strain dominance across a wide variety of carbon sources and strains.

### Strain-specific growth capacities modulate community composition

We next used reconstituted communities to query whether growth properties were completely general to the component strains or were in some cases specific to the strain present in a particular host, and hence could serve as a test of our model. We built a community with the closest relatives of each of the 15 isolates that were available in reference strain collections (Fig. 5A, Methods) and monitored growth of the reference strains in isolation in BHI for one passage and of the community of reference strains for three passages. Grown in isolation, the maximum growth rate of our isolates and their closest-relative reference strains were highly correlated (*R*=0.97, Fig. S17), and the reference community had similar maximum growth rate (Fig. 5B) as the community reconstituted from our isolates. However, the reference community yield was substantially lower than that of the community reconstituted from our 15 isolates or the parent hmSIC (Fig. 5B), likely because *E. fergusonii* had the highest yield of all our isolates but the corresponding reference strain exhibited a substantially lower (>2-fold) yield in BHI (Fig. 5C); all other isolates had similar yield as their reference strain (Fig. 5C). Thus, even if most members do not exhibit relevant strain-level growth differences, the ability of co-cultures to recapitulate a complex community in certain media can be dependent on the individual strain for a few members, especially one of the dominant species.

**Figure 5:**
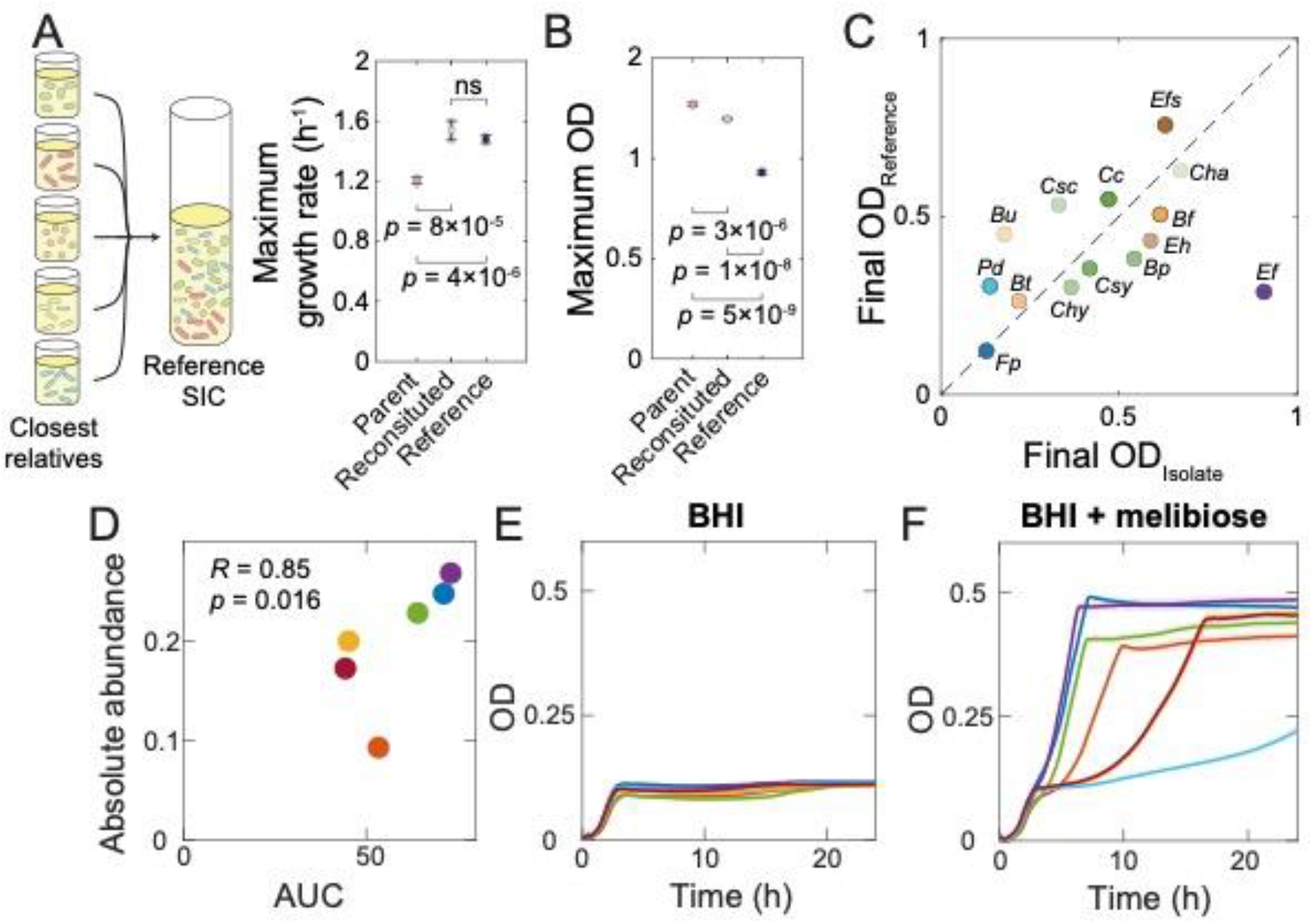
Strain-level growth properties predict community composition. A) Versions of the 15-member SIC were reconstituted with the strains isolated from the hmSIC (“reconstituted SIC”) or their closest relatives from culture collections (“reference SIC”). The maximum growth rate was similar between the reconstituted and reference SIC, whereas the parent (hmSIC) was significantly slower growing. B,C) The yield of the reference SIC was substantially lower than the reconstituted SIC (B), likely due to the lower yield of the reference *E. fergusonii* (*Ef*) strain compared with the isolate from the hmSIC (C). D-F) Across donors, the absolute abundance of the dominant Enterobacteriaceae family ASV was highly correlated with the area under the curve (AUC) of the growth curves of Enterobacteriaceae isolates from each donor in 10% BHI+melibiose (D). The growth of all isolates was similar in 10% BHI (E), whereas each strain displayed distinct dynamics in 10% BHI+melibiose (F).

While in most cases growth in a particular BHI-based medium was similar across SICs from the 8 donors despite the differences in component species (Fig. 1C,1D, S5A,S5B), there were a few cases such as melibiose in which the SICs displayed large variation across donors, particularly in the absolute abundance of Enterobacteriaceae (Fig. 5D), suggesting differences in the metabolic potential of members of this family. We isolated one Enterobacteriaceae from each of the donors (except donor 1); these 7 isolates represented at least 5 distinct 16S sequences, indicating that the similarities in growth of SICs from the 8 donors in most supplemental carbon sources are not due to having the same dominant Enterobacteriaceae strain. The isolates grew similarly in most of the 23 BHI-based media, including base medium only (Fig. 5E). However, in melibiose, their growth curves differed greatly across isolates, and the absolute abundance of each isolate in its corresponding donor community was proportional to the isolate growth curve AUC (Fig. 5D,F). These findings further support that assembly of a community can be understood quantitatively from the growth capacities of its constituent strains.

## Discussion

Our data presents a relatively simple picture of community assembly, in which a natural separation of time scales between the growth of constituent species (Fig. 2B-D) due to differences in growth rate and/or lag time in the base medium allows the fastest grower to have a head start on a supplemental carbon source that is often sufficient for domination. A recent theoretical study predicted that culturing with serial dilution in a nutrient-rich medium (such as BHI) should result in changes to community diversity due to fast-growing species gaining an advantage early on in the batch culturing process (Erez et al., 2020). Our data provide additional nuance to this “early bird” effect, in which sparsity of nutrient consumption and lack of niche overlap (Fig. 3J) prevent the early growers from expanding beyond a well-defined niche (Fig. 3I), which our data predict is approximately conserved for most families across communities (Fig. 2). Will the bacterial early bird always get the carbon worm? In principle, a competitor could make an inhibitory compound, but it would be challenging for the inhibition to have a large impact if all species start at low density due to batch dilution. Regardless, we were able to predict that the early bird would win across a wide range of communities (Fig. 4). In the 11-member community, our model largely predicted the dominance of *C. clostridioforme* (Fig. 4F), indicating that our model is not specific to the fastest growing Enterobacteriaceae and Enterococcaceae. However, certain *Bacteroides* and Ruminococcoaceae species exhibited less growth than expected based on isolated growth measurements, potentially due to the presence of cross-feeding and/or inhibitor compounds that are absent from growth in isolation.

Our study raises many interesting questions about the nature of nutrient utilization and the separation of niches across gut microbial communities. Since the abundance of many species is independent of a supplemental simple sugar despite the increased abundance of *E. fergusonii* and/or Enterococcaceae species, these fast growers are likely not producing a substantial number of metabolites whose cross-feeding potential increases with concentration. The more rapid growth of *Bacteroides* species in a community compared with in isolation (Fig. 2C,3F) may be due to other species synthesizing cofactors such as hemin (Huus et al., 2021). For complex sugars, there may be substantially more potential for cross-feeding as the polysaccharides are broken down; the simple assembly rules that we have elaborated for simple sugars suggest that outliers during growth with a complex sugar may be interpretable as cross-feeding.

The inability of other species to compete for most (if not all) of the nutrients that *E. fergusonii* consumes in BHI, and similarly that *E. fergusonii* largely does not encroach on the niches of other organisms during repeated passaging, may be due to phylum-level metabolic differences. It will be intriguing to determine whether other Proteobacteria have similarly distinct niches from non-Proteobacteria gut commensals, and more generally to quantify the extent of niche overlap among gut commensals. One possibility is that the fastest growers have been selected to consume distinct nutrients; interestingly, there appears to be little niche overlap between *E. fergusonii* and the Enterococcaceae members in our community, some of which can grow almost as quickly as *E. fergusonii* (Fig. 2C). Our data suggest that most species in our community have a niche that is at least partially distinct from the other species, and a consumer-resource model based on this hypothesis of sparse resource consumption quantitatively recapitulated the relationship between OD and richness (Fig. 3J). More generally, the predicted saturation of richness may enable estimation of the number of niches within a particular medium, at least in the context of a starting pool of species.

The ability to dissect the behavior of the parent hmSIC using a reconstituted community from which individual species can be dropped out (Fig. 2A) was critical for testing the “early bird” hypothesis. The 15-member reconstituted community largely recapitulated the growth and composition of the full community (Fig. S16A), likely because it contained most major families, which has been suggested to be the most relevant taxonomic level for grouping similar metabolic capabilities (Estrela et al., 2022b; Goldford *et al*., 2018; Louca et al., 2016; Tian et al., 2020). Our findings suggest that reconstituted communities with relatively tractable numbers of species can serve as reasonable models for more diverse communities. Moreover, since *E. coli* grew similarly to the *E. fergusonii* from our reconstituted community across all media (Fig. 2B), mechanistic studies that disrupt individual genes, for instance using existing genetic libraries (Baba et al., 2006), can easily be carried out in the future with a defined community. On the other hand, complex, undefined communities provide the ability to test for generality, such as the composition across nutrients (Fig. 1), and for functions such as colonization resistance in which the important members are unknown (Aranda-Diaz *et al*., 2022). In the future, a combination of undefined, stool-derived communities with phage editing to remove species would combine the benefits of top-down and bottom-up approaches.

Our findings indicate that isolate growth dynamics are sufficient to predict the identities and even abundances (Fig. 4D) of the dominant members across BHI supplemented with virtually any carbon source from central carbon metabolism. Several factors could affect the generalizability of this conclusion. Our model is based on the observation that all isolates consume nutrients in BHI before transitioning to the supplemented carbon source (Fig. 2C-D). If one or more of the species were to immediately start consuming the carbon source, such behavior would be evident from isolate growth curves, and it remains possible that a different metric that is still based on the “early bird” principle would be able to predict community composition. It will be intriguing to investigate community assembly behavior in other complex media, and our successes in BHI motivate measurements of the growth of more isolates to enable more accurate prediction of assembly dynamics. The potential for strain-specific differences in growth, particularly among the dominant Enterobacteriaceae members (Fig. 5), also motivates studying isolates that derive from the same host. In the future, it may even be possible to predict isolate growth from genomic information, which screens of growth (Tramontano *et al*., 2018) and metabolite consumption and production (Han *et al*., 2021) will facilitate. In total, our findings demonstrate that the assembly of diverse communities can be surprisingly predictable, perhaps as a result of (rather than despite) the inherent complexity of the community and its nutrient environment.

## Methods

### Growth media

For most experiments, we used either a minimal medium with M9 salts or a complex medium consisting of a mixture of M9 salts and 10% Brain Heart Infusion medium (BHI-based medium), each supplemented with a carbon source or ddH_2_O, to culture feces or isolates. Where indicated, we used 100% BHI without M9 salts or supplemental carbon sources. All reagents were pre-reduced by incubation in an anaerobic chamber for at least 24 h.

We prepared 10X supplemental carbon source stock solutions (at 0.7 C-mol/L) by diluting the supplemental carbon sources in H_2_O and sterilizing with a 0.2-μm filter. We used the following supplemental carbon sources: D-lactose (Frontier Scientific, Cat. #502064212); D-glucose (Fisher Chemical, D163); D-galactose (Sigma-Aldrich, Cat. #48260); sucrose (BeanTown Chemical, Cat. #143635); D-fructose (Sigma-Aldrich, Cat. #F0127); L-rhamnose (Sigma-Aldrich, Cat. #83650); glycerol (Fisher BioReagents, Cat. #BP229); D-ribose (Sigma-Aldrich, Cat. #R9629); pyruvic acid, sodium salt (MP Biomedicals, Cat. #194734); oxaloacetic acid (Sigma-Aldrich, Cat. #O7753); sodium citrate, trisodium salt dihydrate (MP Biomedicals, Cat. #194868); D-maltose monohydrate (Sigma-Aldrich, Cat. #M2250); D-melibiose (Sigma-Aldrich, Cat. #63630); D-mannose (Acros Organics, Cat. #150600250); D-mannitol (Acros Organics, Cat.

#1253450000); D-sorbitol (Sigma-Aldrich, Cat. #W302902); galactitol (dulcitol, Acros Organics, Cat. #117700250); L-arabinose (Sigma-Aldrich, Cat. #A3256); L-fucose (Sigma-Aldrich, Cat. #F2252); acetic acid (Fisher Bioreagents, Cat. #BP1185); succinic acid, disodium salt hexahydrate (Acros Organics, Cat. #158751000); and sodium fumarate (Acros Organics, Cat. #215531000). Stock solutions were stored at 4 °C and aliquots (<100 μL in 96-well plates, covered with breathable seals) were pre-reduced in the anaerobic chamber at room temperature for 24-48 h before the start of each experiment.

Before each experiment, stock solutions of the supplemental carbon sources were mixed at 1:10 ratio with 1.1X solutions of M9 and/or M9+BHI. We mixed 5X M9 minimal salts (Sigma-Aldrich, Cat. #M9956), 1M CaCl_2_ (VWR, Cat. #E506) and 1M MgSO_4_ (Boston BioProducts BM-665) to a final concentration of 1X M9 minimal salts, 0.1 mM CaCl_2_, and 2 mM MgSO_4_ with sterile water. For BHI-based media, the solution also contained 10% BHI (BD, Cat. #2237500) that had been autoclaved for 30 min and pre-reduced.

### Passaging of human SICs

Feces from 8 healthy adults were collected and transferred to an anaerobic chamber within 30 min of collection. Approximately 1 g of feces was diluted in 10 mL PBS and vortexed to form a slurry. After 1 h of soaking, the slurry was mixed and allowed to stand for ~5 min so that the largest particles would settle. To initialize passaging, 11.1 μL of the fecal slurry were inoculated into 20 mL of M9 or BHI-based medium without supplemental carbon sources. One hundred eighty microliters of the mixture were aliquoted into shallow 96-well plates and 20 μL of 0.7 C-mol/L supplemental carbon source was added. Each of the 368 conditions (8 human donors, 2 base media, 22 supplemental carbon sources or ddH2O) were inoculated in quadruplicate. The resulting 1,472 SICs were arranged in 16 96-well plates and incubated in the anaerobic chamber at 37 °C (by setting the temperature of the chamber to 37 °C) in a plate stacker (BioTek BioStack 4) with 2 control plates containing uninoculated media at the top and bottom of the stack to minimize edge effects. The stacker was coupled to a Synergy 2 plate reader (BioTek), which read OD and shook the plates for 5 min as often as allowed by the plate reader (~1 h).

As in (Aranda-Diaz *et al*., 2022), plates were incubated for 48 h and then 1 μL of each saturated culture was transferred into 200 μL of the appropriate medium in a new plate, which was covered with a transparent seal poked with ~0.5-mm holes to allow for gas exchange. To minimize contamination and limit the time of experimental manipulation, we used a BenchSmart 96 semi-automatic pipetting system (Rainin) housed in the anaerobic chamber. Each 48-h cycle from inoculation to saturation is referred to as a passage. At the end of passages 1, 3, and 7, a portion of each culture was used to make frozen stocks in 5% DMSO or 25% glycerol, which were stored at −80 °C in sealed bags to reduce exposure to oxygen. The remainder of each culture was stored at −80 °C for DNA extraction and sequencing.

### Isolation of strains

SICs were resuspended in PBS and plated onto BHI, BHI-S, or GAM plates. Plates were incubated at 37 °C in an anaerobic chamber. After 2 days, colonies were grown in the medium corresponding to the plate from which they were isolated for 2 days, and glycerol stocked. The strains were identified via high-throughput MALDI-TOF mass spectrometry of whole colonies with a MALDI Biotyper (Bruker), following manufacturer’s instructions and including formic acid lysis. For the parent hmSIC that we attempted to reconstitute, 15 distinct strains were obtained and their taxonomy was checked by Sanger sequencing. Genomic DNA was extracted from pure cultures using a DNeasy UltraClean 96 Microbial Kit (Qiagen 10196-4) or DNeasy Blood and Tissue Kit (Qiagen 69504) kit. The 16S gene was amplified using primers 5’AGAGTTTGATCCTGGCTCAG and 5’GACGGGCGGTGWGTRCA, and the amplicon was Sanger-sequenced. Taxonomic assignment was performed by alignment using BLAST against the 16S ribosomal RNA sequences (Bacterial and Archaea) database.

To isolate Enterobacteriaceae and Enterococcaceae from human SICs, frozen stocks of the 7^th^ passage in BHI-based medium with supplemental glucose were inoculated into 3 mL BHI (with no supplemental carbon sources) and grown at 37 °C for 48 h. The saturated cultures were serially diluted in PBS and plated onto BHI agar plates. The plates were incubated aerobically at 37 °C for 24 h. Single colonies were struck onto BHI agar plates, from which single colonies were grown in 3 mL BHI for 24 h and made into glycerol stocks. To identify the strains, DNA was extracted from cultures using a DNeasy UltraClean 96 Microbial Kit (Qiagen, Cat. #10196-4). The 16S gene was amplified using primers 5’AGAGTTTGATCCTGGCTCAG and 5’GACGGGCGGTGWGTRCA, and the amplicon was Sanger-sequenced. Taxonomic assignment was performed by alignment using BLAST against the 16S ribosomal RNA sequences (Bacterial and Archaea) database (as of 9/28/2020).

### Community reconstitution

For community reconstitution, frozen stocks of isolated (Table S1) or reference (Table S2) strains were streaked onto BHI 5% sheep blood agar plates and incubated for 48 h at 37 °C in anaerobic conditions. A single colony of each strain was inoculated into 3 mL of BHI in isolation, and incubated for 48 h at 37 °C. The OD was measured for each strain and diluted to a final OD of 10^-4^ before mixing in 200 μL of BHI in a 96-well plate. The mixed cultures were grown for 48 h in a plate reader to measure growth. After 48 h, the cultures were diluted 1:200 into fresh BHI.

### Growth measurements

To measure the growth of SICs or individual strains, they were first grown from a frozen stock in BHI to effectively remove the glycerol in the frozen stocks. For SICs, approximately 10 μL of frozen stock were inoculated in 3 mL BHI and grown at 37 °C for 48 h for 2 passages, and diluted 1:200 after the first passage. For individual strains, frozen stocks were struck for single colonies onto BHI-blood or BHI agar plates and incubated at 37 °C for 48 h. A single colony, or a group of colonies for strains that grew slowly, was inoculated in 3 mL of BHI and grown at 37 °C for 48 h. For isolate co-cultures (reconstituted SICs), equal volumes of the isolate cultures were mixed in a total of 1 mL, diluted 1:200 into 3 mL of BHI, and grown for 48 h at 37 °C. The SIC, isolate, or reconstituted SIC cultures were then diluted 1:200 into four replicates in 96-well microplates in media as described above and grown in an Epoch2 or Synergy 2 plate reader (Biotek) for 48 h with constant shaking. SICs were further passaged after 48 h by diluting 1:200 into the corresponding fresh media. SIC cultures were collected at the end of each passage and stored at −80 °C for DNA extraction.

### 16S sequencing

DNA was extracted from cultures using the DNeasy UltraClean 96 Microbial Kit (Qiagen, Cat. #10196-4). The bacterial 16S rRNA variable region 4 (V4) was amplified using Earth Microbiome Project-recommended 515F/806R primer pairs using the 5PRIME HotMasterMix (Quantabio, Cat. #2200410) with the following program in a thermocycler: 94 °C for 3 min, 35 cycles of [94 °C for 45 s, 50 °C for 60 s, and 72 °C for 90 s], followed by 72 °C for 10 min. PCR products were cleaned with the UltraClean 96 PCR Cleanup kit (Qiagen, Cat. #12596-4) and pooled using the same volume for each sample. Pooled libraries were concentrated by ethanol precipitation and purified by gel extraction of the corresponding library size using the NucleoSpin Gel and PCR Clean-up Mini kit (Macherey-Nagel). Libraries were sequenced using the MiSeq Reagent Kit v3 with 300-bp paired end reads on a MiSeq (Illumina).

Demultiplexed fastq files for each sample were processed using DADA2 (Callahan et al., 2016) with the following parameters for the `filterAndTrim` function: `truncLenF = 250, truncLenR = 180, maxEE = c(2,2), truncQ = 2, maxN = 0`. Default parameters were used for the `learnErrors` and `dada` functions. Taxonomic assignment was performed with the `assignTaxonomy` function using the Greengene Database (gg_13_8_train_set_97.fa.gz).

### Sampling of cultures during a single passage

We serially passaged the hmSIC in BHI-base medium with ddH_2_O or each of the 22 supplemental carbon sources under the growth conditions described above. After 2 passages in 10% BHI in a test tube, we inoculated each culture 1:200 into a flat-bottomed shallow 96-well plate with 200 μL of the appropriate medium (16 replicate plates in total, one per collection time point). At each time point (every 30 min or 1 h for the first 8 h and every 2-16 h thereafter), we collected a plate and stored it at −80 °C until processing for 16S rRNA gene sequencing to quantify community composition.

## QUANTIFICATION AND STATISTICAL ANALYSIS

### 16S data analysis

Custom MATLAB R2018a (MathWorks) scripts were used for 16S data analysis, as in (Aranda-Diaz *et al*., 2022; Celis et al., 2022). In brief, alpha diversity (richness) was measured by rarefying all samples to 5,000 reads and calculating the number of taxa with relative abundance >0. Weighted and unweighted Unifrac distances between samples for beta diversity were calculated using custom MATLAB code. Unifrac was calculated as described in (Lozupone and Knight, 2005). Weighted Unifrac was calculated as described in (Lozupone et al., 2007): for samples *A* and *B*, 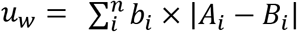, where *n* is the total number of branches in the tree, *b_i_* is the length of branch *i*, and *A_i_* and *B_i_* are the relative abundances of the taxa that descend from branch *i* in samples *A* and *B*, respectively. Only taxa present at >0.01% in more than two samples were used to calculate distances. Principal coordinate analyses were performed on distances between samples with >5,000 reads.

### Absolute abundance calculations

As a proxy for the absolute abundance of a given taxon in a community, we multiplied the taxon’s relative abundance (as measured from 16S sequencing) by SIC yield (OD).

### Growth curve analyses

Maximum growth rate was calculated as the maximum slope of ln(OD) with respect to time (calculated from a linear regression of a sliding window of 11 time points) using custom Matlab R2018b (Mathworks) code. To compare across media, instantaneous growth rates (defined by *d* ln(OD)/*dt*) were normalized to the maximum value.

### Consumer-resource simulations and sparsity model

We considered a consumer-resource model with *N* consumers and *M* resources, in which 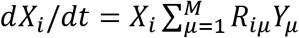 and 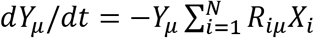. Here, *X_i_* is the absolute abundance of consumer *i, Y_μ_* is the amount of resource *μ*, and *R_iμ_* is the consumption rate of resource *μ* by consumer *i*. Within this model, all resources that can be consumed will be depleted and converted to consumer abundances at large times, regardless of initial conditions or consumption rates. We also describe the niche overlap among consumers by a sparsity parameter *S*, the probability that a consumer does not consume a resource. In this scenario, in the limit of large *M* (as is reasonable for a rich medium like BHI), the fraction of resources in a community with richness *d* that is not consumed is *S^d^*, hence the fraction of resources consumed and converted to total consumer abundance is 1 – *S^d^*. This relation was fitted to determine S from experimental data of normalized OD (fraction of resources consumed) and richness *(d)* by minimizing the mean squared error between observed and predicted normalized OD.

### Flux Balance Analysis

We used Flux Balance Analysis (FBA) to quantify the maximum theoretical biomass yield obtainable when consuming one of the 22 carbon sources used in our experiments. For this analysis, we used iJO1366, a high-quality, genome-scale model of the *E. coli* metabolic network (Orth et al., 2011). We added to iJO1366 the oxaloacetate exchange and transporter reactions (EX_oaa_e, OAAt2_2pp, OAAtex) to allow for growth on oxaloacetate (by default the iJO1366 model does not include the ability to grow on oxaloacetate). As in previous work (Estrela et al., 2021), we set a lower bound on ATPM to 0 mmol/gDWh and assumed all inorganic compounds (ca2_e, cbl1_e, cl_e, co2_e, cobalt2_e, cu2_e, fe2_e, fe3_e, h_e, h2o_e, k_e, mg2_e, mn2_e, mobd_e, na1_e, nh4_e, ni2_e, pi_e, sel_e, slnt_e, so4_e and tungs_e, zn2_e) were in excess (a lower bound on the exchange reactions of −1000 mmol/gDWh). A lower bound on the oxygen exchange reaction was set to 0 to reflect the anaerobic conditions of our experiments. To estimate optimal biomass yield, the exchange flux was set to −1 cmol/gDWh for each carbon source and FBA was performed using the biomass reaction as the objective function (BIOMASS_Ec_iJO1366_core_53p95M). This analysis was performed using the COBRAPY package (Ebrahim et al., 2013).

## Author Contributions

A.A.-D, S.E., A.S., and K.C.H designed the research; A.A.-D., S.E., L.W., P.-Y.H., J.V., T.T., T.C., R.Y., T.N., and F.B.Y. performed the research; A.A.-D., S.E., L.W., P.-Y.H., J.V. T.N., A.S., and K.C.H. analyzed the data; and A.A.-D, S.E., L.W., A.S., and K.C.H wrote the paper and all authors reviewed it before submission.

## Acknowledgements

We would like to thank members of the Huang lab for helpful discussions. A.A.-D. is a Howard Hughes Medical Institute International Student Research fellow, a Stanford Bio-X Bowes fellow, and a Siebel Scholar. The authors acknowledge funding from the Allen Discovery Center at Stanford on Systems Modeling of Infection (to K.C.H), NIH Awards R01 AI147023 and RM1 GM135102 (to K.C.H.), and NSF grants EF-2125383 and IOS-2032985 (to K.C.H.). K.C.H. is a Chan Zuckerberg Biohub Investigator.

